# “polishCLR: a Nextflow workflow for polishing PacBio CLR genome assemblies”

**DOI:** 10.1101/2022.02.10.480011

**Authors:** Jennifer Chang, Amanda R. Stahlke, Sivanandan Chudalayandi, Benjamin D. Rosen, Anna K. Childers, Andrew Severin

**Author notes:** co-first authors contributed equally (Jennifer Chang, Amanda R. Stahlke). Author for Correspondence: Andrew Severin, Genome Informatics Facility, Iowa State University, Ames, Iowa, USA, 50010.

## Abstract

Long-read sequencing has revolutionized genome assembly, yielding highly contiguous, chromosome-level contigs. However, assemblies from some third generation long read technologies, such as Pacific Biosciences (PacBio) Continuous Long Reads (CLR), have a high error rate. Such errors can be corrected with short reads through a process called polishing. Although best practices for polishing non-model *de novo* genome assemblies were recently described by the Vertebrate Genome Project (VGP) Assembly community, there is a need for a publicly available, reproducible workflow that can be easily implemented and run on a conventional high performance computing environment. Here, we describe polishCLR (https://github.com/isugifNF/polishCLR), a reproducible Nextflow workflow that implements best practices for polishing assemblies made from CLR data. PolishCLR can be initiated from several input options that extend best practices to suboptimal cases. It also provides re-entry points throughout several key processes including identifying duplicate haplotypes in purge_dups, allowing a break for scaffolding if data are available, and throughout multiple rounds of polishing and evaluation with Arrow and FreeBayes. PolishCLR is containerized and publicly available for the greater assembly community as a tool to complete assemblies from existing, error-prone long-read data.

## Main

Long reads, including those generated by third-generation sequencing platforms such as Pacific Biosciences (PacBio) and Oxford Nanopore Technology (ONT), have revolutionized genome assembly (Childers et al., 2021; Hotaling et al., 2021; Rhie et al., 2021). However, until recent advances (Hon et al., 2020; Wang et al., 2021), long-read sequencing technologies have had high error rates (5-15%), especially among indels (Watson & Warr, 2019). Thus, the vast majority of long-read data that is currently publicly available yields assemblies with low overall consensus accuracy, which, if left uncorrected, negatively impacts downstream analyses, such as gene annotation (Watson & Warr, 2019). These assembly errors require correction with an additional higher fidelity read set, such as short-read Illumina data, in a process called polishing (Chin et al., 2016; Helper et al., 2016; Walker et al., 2014).

Polishing can be a complex process, with high computational cost, non-trivial file-handling, and issues around special cases that must be resolved. For example, the long-read contig assembly should ideally be polished with high-fidelity reads from the same individual, but this may not be technically feasible when sufficient DNA cannot be extracted from individual specimens, for example in small-bodied organisms such as many insects. In such cases, it is necessary to modify parameters in the standard workflow. Best practices for *de novo*, chromosome-scale vertebrate genome assembly from error prone PacBio continuous long reads (CLR) reads were recently described (Rhie et al., 2021), however it can be challenging to run this code and reproduce widely. In order to produce the best possible genome assemblies using existing data from species regardless of their position in the tree of life, the genome assembly community needs a publicly available, flexible and reproducible workflow that is containerized so it can be run on any conventional HPC.

Bioinformatic pipelines with complex entrance and decision points, such as polishing, are inherently difficult to track, develop, and debug. Increasing interest in workflow development systems that track data and software provenance, enable scalability and reproducibility, and re-entrant code (Wratten et al., 2021) have led to the development of several workflow languages, largely inspired by GNU Make (Amstutz et al., 2016; Köster & Rahmann, 2012; Stallman & McGrath, 1991). Nextflow is a Domain Specific Language (Di Tommaso et al., 2017) that currently leads workflow systems in terms of ease of scripting and submitting to cloud computing resources (Fjukstad & Bongo, 2017; Jackson et al., 2021; Leipzig, 2017; Spjuth et al., 2020). A key benefit of Nextflow compared to earlier workflow languages is being able to submit jobs to a local machine, an HPC, or cloud-based compute environments. These features empower a large range of bioinformatic pipelines, for example, initial read processing and annotation lift-over (Federico et al., 2019; Talenti & Prendergast, 2021). In this paper, we describe polishCLR, a reproducible Nextflow workflow which implements the current best practices for polishing CLR assemblies and is flexible to multiple input assembly and sample considerations.

The polishCLR workflow can be easily initiated from three input cases (Fig. 1). In the first case (Case 1), users may start with an unresolved primary assembly with (e.g., the output of FALCON 2-asm (Chin et al., 2016)) or without (e.g., the output of Canu or wtdbg2 (Koren et al., 2017; Ruan & Li, 2020)) associated contigs. Additionally, it can handle a haplotype-resolved but unpolished set (Case 2) (e.g., the output of FALCON-Unzip 3-unzip (Chin et al., 2016)). In the ideal case (Case 3), the pipeline is initiated with a haplotype-resolved, CLR long-read polished set of primary and alternate contigs (e.g., the output of FALCON-Unzip 4-polish). In all cases, inclusion of organellar genomes, e.g., the mitochondrial genome, will improve the polishing of nuclear mitochondrial or plasmid pseudogenes (Howe et al., 2021). Organellar genomes should be generated and polished separately for best results, using pipelines such as the mitochondrial companion to polishCLR, polishCLRmt (Stahlke et al, in prep) or mitoVGP (Formenti et al., 2021). To allow for the inclusion of scaffolding before final polishing (e.g., Durand et al., 2016) and increase the potential for gap-filling across correctly oriented scaffolded contigs, the core workflow is divided into two steps, controlled by a --step parameter flag.

**Figure 1.**
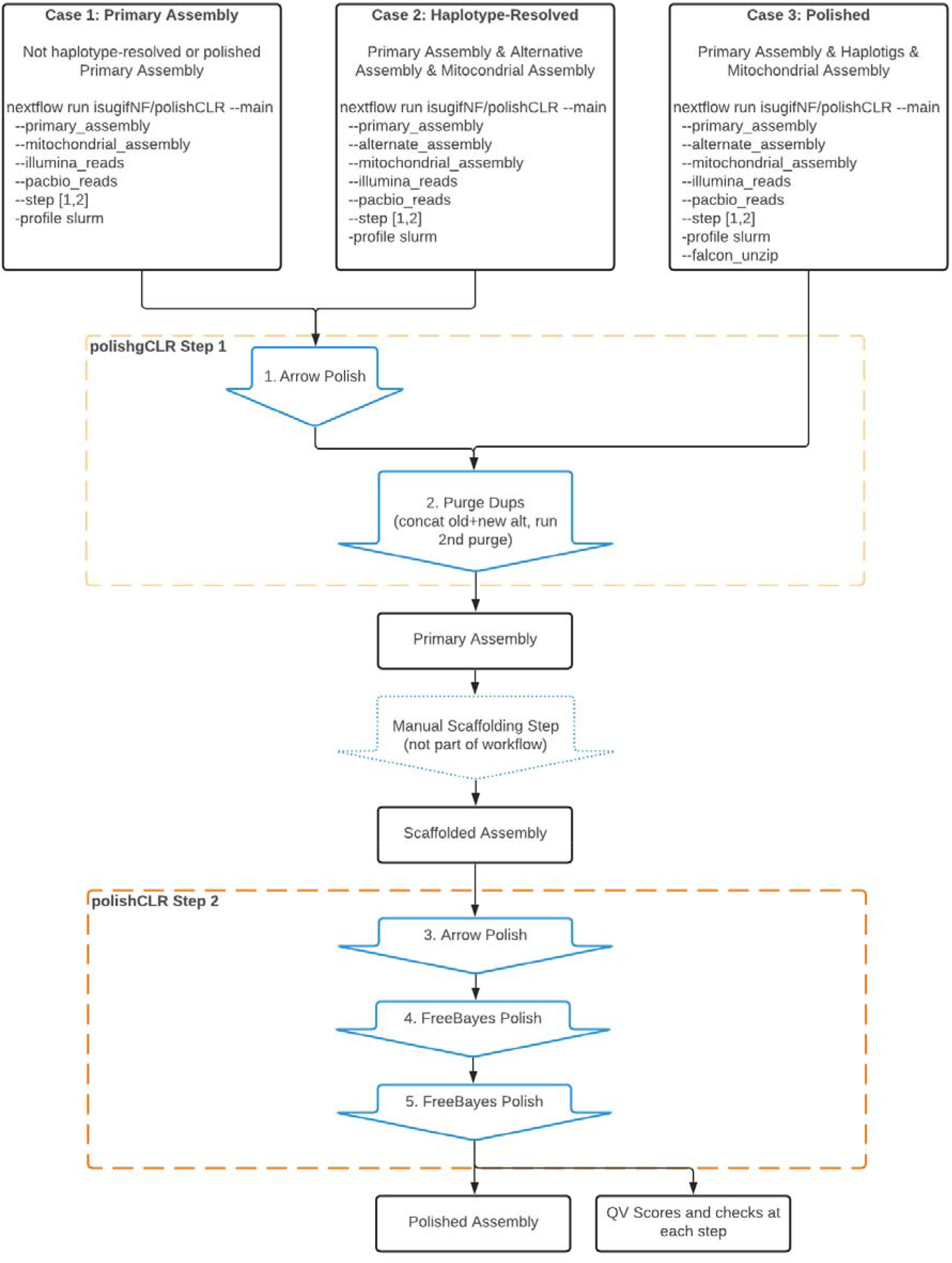
Diagram of polishCLR workflow for three input cases. Polishing steps 1 and 2 are run separately to accommodate an optional scaffolding step. Blue arrows indicate processes while black boxes indicate products. The dotted arrow indicates that the manual scaffolding step is optional and not within the scope of this pipeline.

In --step 1, if initiating the workflow under Case 1 or 2, unpolished primary contigs are merged with the organellar genome and associated contigs or alternate haplotypes if available, then polished with a single round of Arrow long-read polishing (Pacific BioScience) before entering the core workflow (controlled by --arrow01 true). During Arrow steps (here and later in Step 2), polishCLR improves re-entry and computational resource use by delineating at least seven Nextflow processes: 1) indexing each contig, 2) creating a pbmm2 index of the assembly, 3) aligning PacBio reads to the assembly, 4) submitting a GCpp Arrow job for each contig in parallel, 5) combining the separate contig variant calling format (VCF) files, 6) reformatting Arrow generated VCF for Merfin filtering (Formenti et al., 2021), and 7) converting the resultant VCF back to FASTA format. Then, in all three cases, the core workflow employs purge_dups v.1.2.5 (Guan et al., 2020) to remove duplicated sequence at the ends of separated primary contigs, with cutoffs automatically estimated from a generated histogram of k-mers. The histogram is captured as one of the relevant outputs for users to review. Purged primary sequences are then concatenated to the alternate haplotype contigs and the combined alternate set is purged of duplicates. BUSCO completeness metrics (Manni et al., 2021; Seppey et al., 2019; Simao et al., 2015; Waterhouse et al., 2018) are generated for the primary contigs before and after removing duplicated content to ensure that cutoff parameters are effective and do not remove too much genic content. The eukaryotic BUSCO database is used by default, but users may provide a designated lineage (controlled by a --busco-lineage flag). If additional data are available, this de-duplicated primary contig assembly can then be scaffolded by the user before initiating the second phase of the workflow.

In --step 2, the primary, alternate, and organellar assemblies are merged and polished with an additional round of Arrow, followed by two rounds of FreeBayes (Garrison & Marth, 2012). Indeed, the iterative nature of polishing benefits from the re-entrant caching and templates of the workflow. By default, this second round of Arrow-identified variants are only filtered via Merfin if the CLR and the Illumina reads came from the same specimen, adding additional Nextflow processes to the Arrow delineation described above to create a meryl genome database and perform filtering (McCartney et. al, unpublished data). However, if short-read data are from a different specimen than the long-read-based contig assembly, then Merfin filtering can be turned off to avoid over-polishing with the parameter flag --same-specimen false. As with Arrow, polishCLR takes advantage of Nextflow in seven processes to implement FreeBayes: 1) creating contig windows, 2) generating a meryl database from the genome, 3) aligning Illumina short reads, 4) polishing via FreeBayes, 5) combining windowed VCF files, 6) filtering VCFs by Merfin, 7) and converting VCFs to FASTA. Throughout the polishCLR workflow, reports are automatically generated to assess genome assembly quality, including k-mer based completeness and consensus accuracy QV scores via Merqury (Rhie et al., 2020), as well as genome size distribution statistics generated with BBMap (e.g., N50) (Bushnell, 2014). These reports allow users to understand how the assembly changed through each major phase of the workflow. The complete, detailed pipeline can be viewed in Supplemental Figure 1.

The polishCLR workflow is publicly available (https://github.com/isugifNF/polishCLR), reproducible, interoperable, easily portable, and can be run on a conventional HPC or extended to cloud computing resources simply by swapping out the Nextflow config file. Software dependencies are listed in a conda environment file. Its use has been demonstrated on several arthropod species assemblies as part of the Ag100Pest Initiative (Childers et al., 2021). Runtimes and summaries from each of the three starting input cases are included (Supplementary Table S1; Stahlke and Coates, 2022) with a full genome announcement forthcoming (Stahlke et al., unpublished data). The polishCLR workflow will increase the efficiency of polishing many genomes and reduce the potential of human error in this multistep process. Despite the much-reduced error rate of PacBio HiFi and ONT reads, polishing approaches continue to be an important component of accurate genome assembly (Shafin et al., 2021). Although this pipeline was not designed to polish with ONT reads, the workflow is available on GitHub and welcomes any future contributions.

## Acknowledgements

This work was supported by the U.S. Department of Agriculture, Agricultural Research Service (USDA-ARS) and used resources provided by the SCINet project of the USDA-ARS, ARS project number 0500-00093-001-00-D. JC was supported, in part, by an appointment to the Research Participation Program at the Agricultural Research Service, United States Department of Agriculture, administered by the Oak Ridge Institute for Science and Education through an interagency agreement between the U.S. Department of Energy and USDA-ARS under contract number DE-AC05-06OR23100. This workflow was developed as part of the USDA-ARS Ag100Pest Initiative. The authors thank members of the USDA-ARS Ag100Pest Team and SCINet Virtual Resource Support Core (VRSC) for fruitful discussions and troubleshooting throughout the development of this workflow. All opinions expressed in this paper are the author's and do not necessarily reflect the policies and views of USDA, DOE, or ORAU/ORISE. Mention of trade names or commercial products in this publication is solely for the purpose of providing specific information and does not imply recommendation or endorsement by the U.S. Department of Agriculture. USDA is an equal opportunity provider and employer.

## Figures & Tables

**Supplemental Figure 1.**
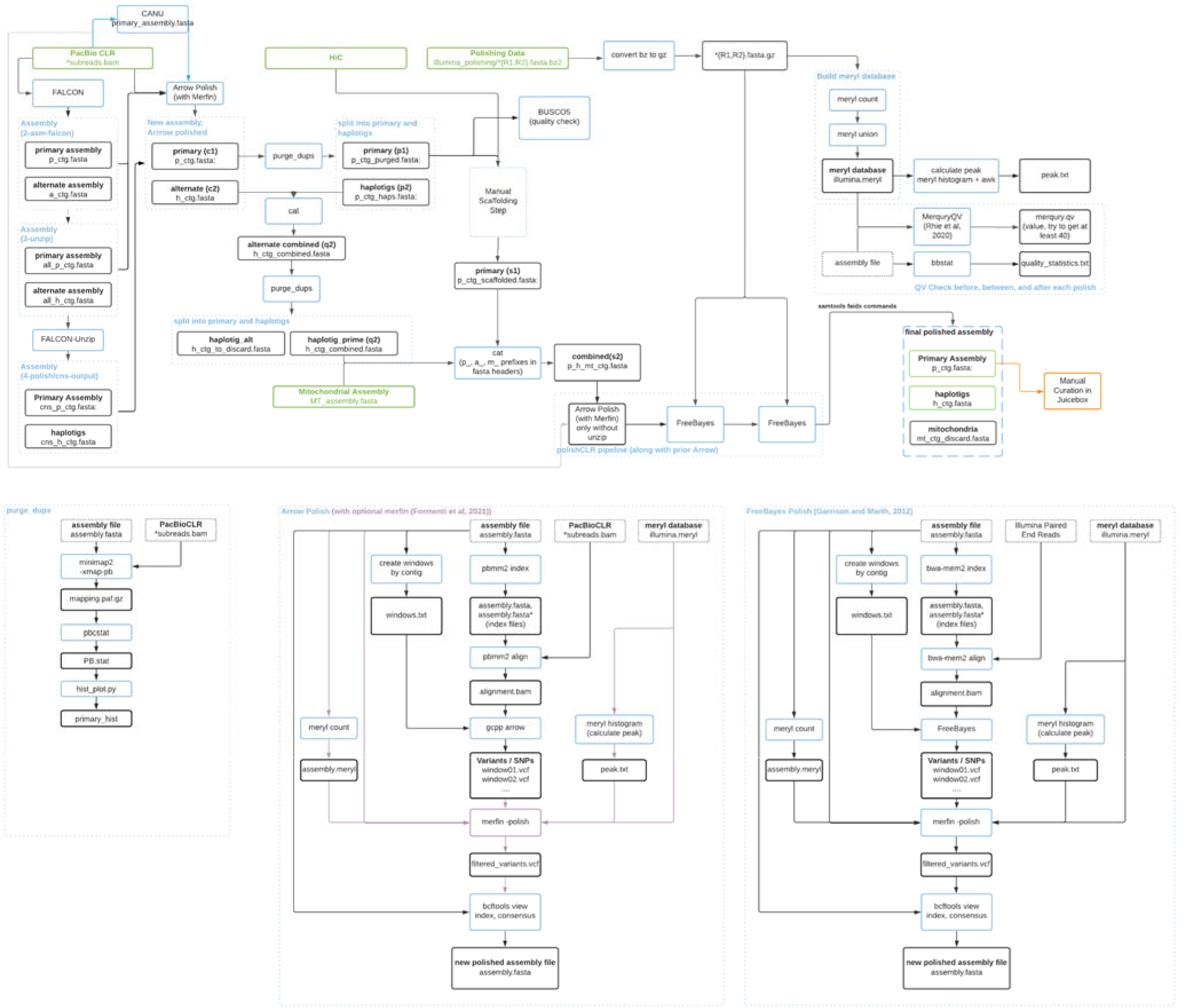
Detailed diagram of the polishCLR pipeline. PacBio CLR long reads, Hi-C data, Illumina short reads, and organellar input data are shown in the green boxes. A detailed view of the Arrow and FreeBayes polish steps are expanded below separately, with the optional Merfin filtering during the Arrow polishing step shown in purple. Merfin filtering is part of all FreeBayes polishing steps.

**Supplemental Table 1.** The polishCLR workflow was benchmarked on the primary contigs of *Helicoverpa zea* generated by FALCON (Chin et al. 2016). Metrics for each assembly include starting pseudo-haploid primary and alternate combined genome size (Mb) and number of contigs, initial quality scores, CPU hours through the pipeline final quality scores, and final genome size and number of contigs. This table provides an indication of scalability of the pipeline on a SLURM managed HPC.

| Case | Input stage of Falcon assembly | Input Genome Size (Mb) / Number of contigs | Starting QV | Final QV | CPU hours | Output Genome Size (Mb) / Number of contigs |
| --- | --- | --- | --- | --- | --- | --- |
| 1 | 2-asm-falcon/ | 501.287 / 789 | 31.8218 | 40.3033 | 211.3 | 500.578 / 799 |
| 2 | 3-unzip/ | 515.499 / 1125 | 31.8492 | 38.9997 | 224.2 | 511.878 / 1022 |
| 3 | 4-polish/ | 509.063 / 882 | 38.8556 | 41.9163 | 195.0 | 509.052 / 882 |

## Notes

https://data.nal.usda.gov/dataset/data-polishclr-example-input-genome-assemblies

https://github.com/isugifNF/polishCLR

### Competing Interest Statement

The authors have declared no competing interest.

