## Supplementary figures and images for "“polishCLR: a Nextflow workflow for polishing PacBio CLR genome assemblies”"

### Supplemental Figure 1

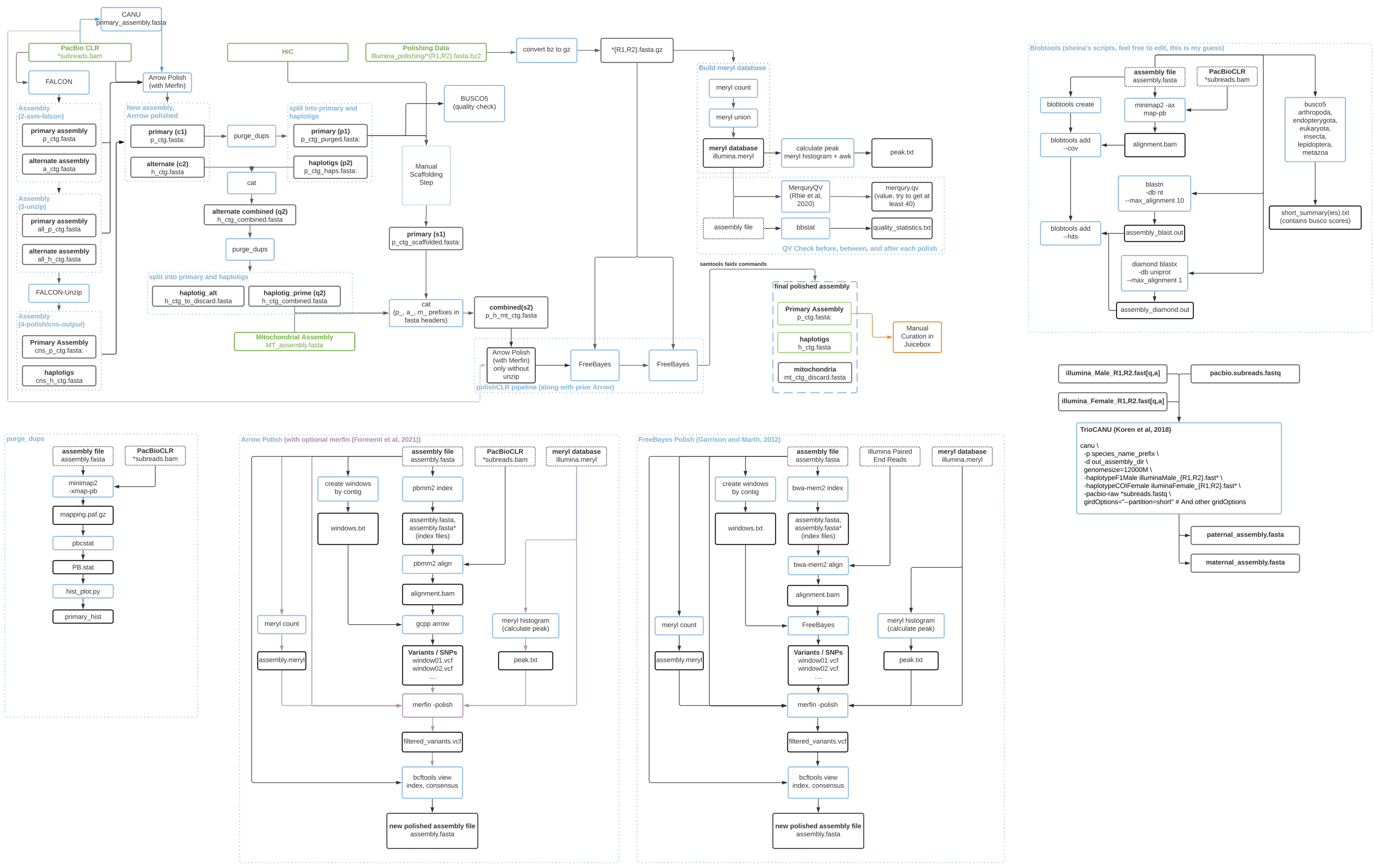
